# PathExpSurv: Pathway Expansion for Explainable Survival Analysis and Disease Gene Discovery

**DOI:** 10.1101/2022.11.08.515625

**Authors:** Zhichao Hou, Jiacheng Leng, Jiating Yu, Zheng Xia, Ling-Yun Wu

## Abstract

**Motivation:** In the field of biology and medicine, the interpretability and accuracy are both important when designing predictive models. The interpretability of many machine learning models such as neural networks is still a challenge. Recently, many researchers utilized prior information such as biological pathways to develop bioinformatics methods based on neural networks, so that the prior information can provide some insights and interpretability for the models. However, the prior biological knowledge may be incomplete and there still exists some unknown information to be explored.

**Results:** We proposed a novel method, named PathExpSurv, to gain an insight into the black-box model of neural network for cancer survival analysis. We demonstrated that PathExpSurv could not only incorporate the known prior information into the model, but also explore the unknown possible expansion to the existing pathways. We performed downstream analyses based on the expanded pathways and successfully identified some key genes associated with the diseases and original pathways.

**Availability:** Python source code of PathExpSurv is freely available at https://github.com/Wu-Lab/PathExpSurv.

**Supplementary information:** Supplementary data are available at *Bioinformatics* online.

## 1 Introduction

When developing a predictive model in the area of biology and medicine, it is significant to balance the trade-off between accuracy and interpretability. Simple models like linear regression usually have high interpretability but don’t perform well, whereas the complex models based on deep learning can achieve good performance but it is hard to explain the black-box inside these models.

There are many different kinds of predictive tasks which can be roughly categorized into classification and regression task. In this paper, we focus on a special regression task, the survival regression, which is developed for dealing with censored data. Survival models are applied to perform time-to-event analysis in order to understand the relationships between the patients’ covariates and the risk of the event. The Cox proportional hazards model (CPH) (Cox, 1972), a semi-parametric regression model, was widely used in survival analysis. This model assumes that the log-risk of failure is a linear combination of the patient’s features. Although linear model has good interpretability, it might be too simplistic to just assume that the log-risk function is linear.

With the advent of machine learning, biomedical researchers were able to fit survival data with more sophisticated nonlinear log-risk functions. Faraggi and Simon (1995) firstly incorporated the feed-forward neural network into Cox proportional hazards model (CPH), but this model with only a single hidden layer hadn’t showed great improvements beyond the CPH. DeepSurv (*Katzman et al*., 2018) was an addition to Simon-Farragi’s network and configurable with multiple hidden layers. It employed a more complex deep neural network to model the relationships between the observed features and the patients’ risk of failure and showed improvements on the CPH when modeling the non-linear data. These neural network-based methods have high predictive performance, but they only leverage the fully connected neural networks, which are arbitrarily over-parameterized and lack of interpretability.

In order to design a biologically informed and sparse neural network, DeepOmix (Zhao *et al*., 2021) utilized signaling pathways as the functional modules based on KEGG and Reactome databases to construct pathway-associated sparse network. Each node encoded some biological entity and each edge represented a known relationship between the corresponding entities. However, this model only considered the known and fixed functional modules in databases to design a sparse network, which might leave out some important factors. In fact, despite painstaking and manual curation, signaling pathways stored in databases still remained incomplete (Ritz *et al*., 2016).

Therefore, it is necessary to make an exploration on the unknown space out of the prior information and identify some significant genes which may complement the original functional modules. In this paper, we presented PathExpSurv, a novel survival analysis method by exploiting and expanding the existing pathways. We firstly incorporated prior biological knowledge of signaling pathways into the neural network for survival analysis. In order to explore the possible unknown pathways with better performance, we further added the genes beyond the databases into the neural network pre-trained using the existing pathways, and continued to train a regularized survival analysis model, with a *L*_1_ penalty that guarantees the sparse structure in the expanded pathways. By simultaneously exploiting the existing pathways and exploring the unknown pathways, PathExpSurv can gain an insight into the black-box model of neural network for survival analysis. We performed some downstream analyses based on the expanded pathways and successfully identified some key genes associated with the diseases and original pathways.

## 2 Methods

### 2.1 Basic Architecture

Suppose *G* is the number of genes, and *N* is the number of samples (patients). PathExpSurv uses a biologically informed neural network *f*_W_(**x**) to predict the effects of a patient’s covariates on their hazard rate, with the input of gene expression 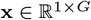 and the learnable weights **W**. Our main objective is to optimize the mean negative log partial likelihood:

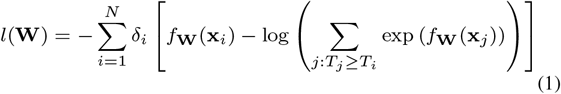

where *δ_i_* ∈ {0, 1} is the event indicator of *i*-th sample, 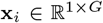 is the feature vector, and 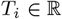 is the event time.

The basic architecture of neural network *f*_W_ (**x**) consists of 3 layers (**Fig**. 1a). The first layer is gene layer, the second layer is pathway layer and the third layer is the output layer. The nodes of first and second layers encode the genes and pathways respectively, and each edge represents the relationship between a gene and a pathway. The connections between the corresponding entities follow the pathway database such as KEGG and are encoded by a mask matrix **M**. We assume that the genes belonging to the same pathway have similar functions, so we constrain the weight **W**_1_ between the gene and pathway layer to be non-negative. The output of neural network is calculated as:

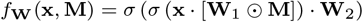

where ⊙ is the element-wise multiplication of two matrices, 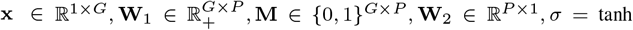, and *P* is the number of pathways explored in the model.

**Fig. 1.**
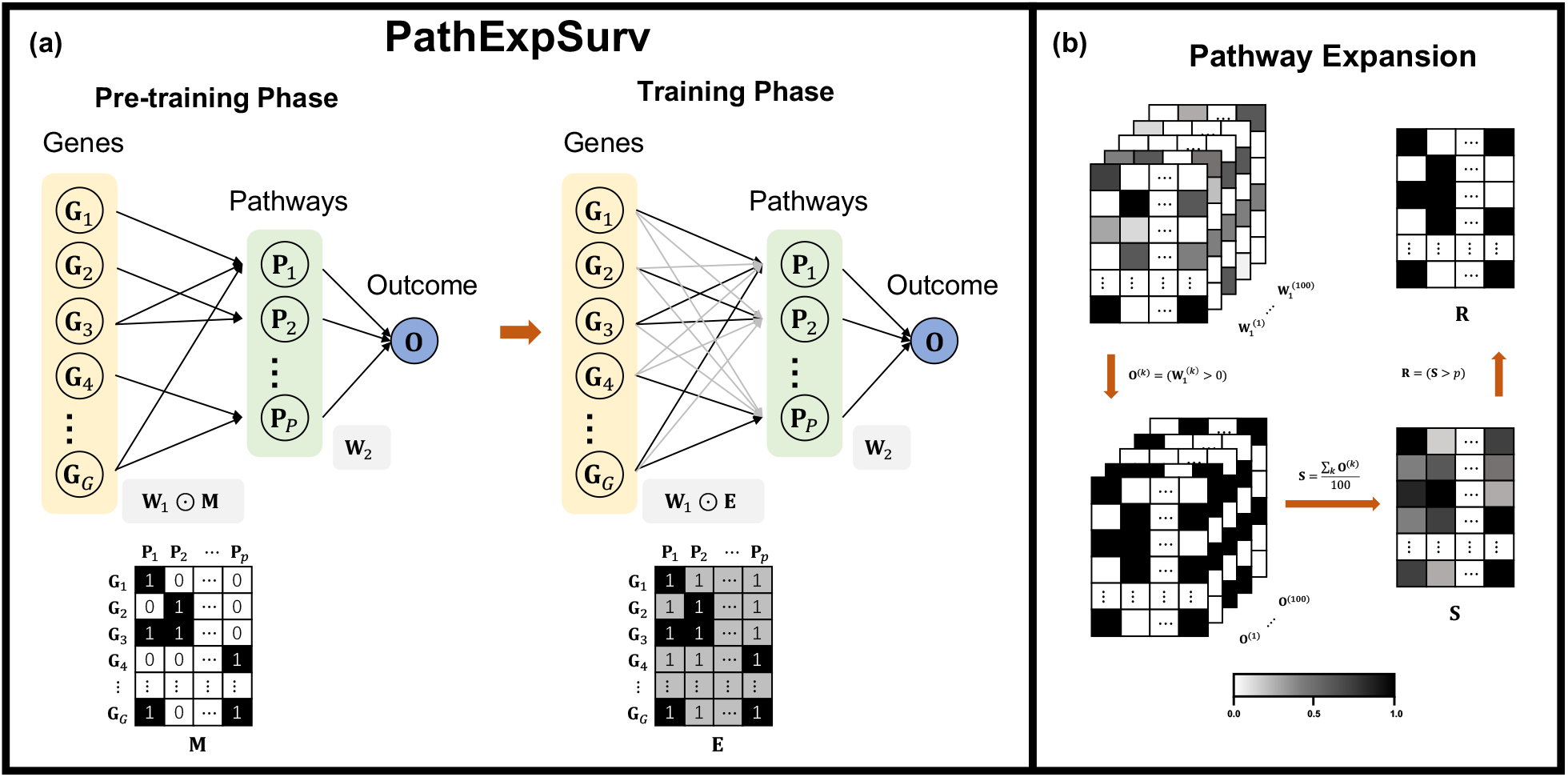
(a) Schematic overview of PathExpSurv. The basic architecture of the neural network consists 3 layers (gene layer, pathway layer and output layer). The connection between the gene layer and the pathway layer is determined by the pathway mask matrix, in which number 1 (black) means a non-penalized link representing a fixed relationship between gene and pathway in prior information, number 1 (grey) means a penalized link representing a possible relationship to be explored, and number 0 (white) means no link. The training scheme of PathExpSurv includes two phases, namely pre-training phase and training phase. In the pre-training phase, the prior pathway mask (**M**) is used to pre-train the model to achieve a relatively high and stable performance. In the training phase, a specific fully connected mask (**E**) with prior links and *L*_1_-penalized non-prior links is used to train the model to explore the unknown space and obtain the expanded pathways. (b) Pipeline of pathway expansion. We first randomly chose 90% samples from the dataset to train the PathExpSurv model, and repeated 100 times to obtain the weight matrices between the gene layer and the pathway layer 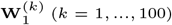. Then we transformed these matrices into binary matrices **O**^(*k*)^ (*k* = 1,…, 100), and calculated the occurrence probability matrix **S** based on these binary matrices. Finally we obtained the expanded pathways matrix **R** by filtering out the gene-pathway pairs with small occurrence probabilities.

### 2.2 Two-Phase Training Scheme

We proposed a novel optimization scheme consisting 2 phases (**Fig.** 1a): pre-training phase and training phase, in order to improve the performance of neural network by expanding the prior pathways.

During the pre-training phase, we utilized the prior pathways from the KEGG database to pre-train the model. We added a standard deviation term to the loss function due to the assumption that the genes in the prior functional modules are almost equally important. Then the objective function of pre-train phase became:

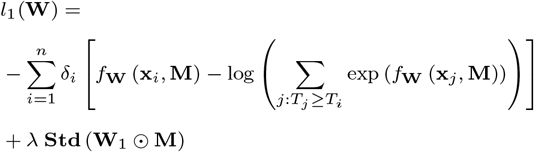

where **M** was the prior pathway mask matrix obtained from the KEGG database.

During the training phase, we changed the connections between the gene layer and the pathway layer to fully connected, and added a *L*_1_ regularization term in order to select a few important genes from the genes outside the prior pathways. That is, we optimized the following loss:

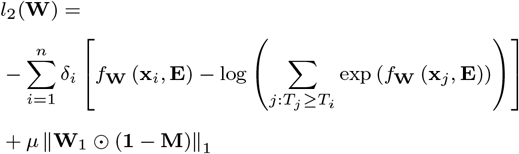

where **E** ∈ {1}^*G* × *P*^ is the matrix of which the elements are all 1.

### 2.3 Evaluation Metric

When performing the evaluation of survival analysis, we need to consider the censored data. The concordance index (C-index) (Uno *et al*., 2011) is the most widely used evaluation metric in survival analysis. C-index is defined as:

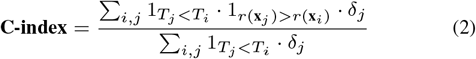

C-index expresses the proportion of concordant pairs in the dataset which estimates the probability that, for a random pair of individuals, the ordering of the predicted hazard risk of the two individuals is concordant with that of their true survival time.

### 2.4 Pathway Expansion

In order to identify the reliable genes complement to the prior pathways, we performed the following procedure as shown in **Fig.** 1b. Firstly, we randomly permuted the dataset and selected 90% samples from the dataset each time to train the PathExpSurv model. In this way, we repeated 100 times and obtained 100 different weight matrices between the gene layer and the pathway layer, 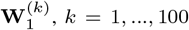. Then we calculated the corresponding occurrence matrix as follows:

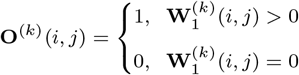

where *k* = 1,…, 100, *i* = 1,…, *G*, *j* = 1,…, *P*.

Secondly, we defined the occurrence probability of *i*-th gene in the *j*-th pathway as:

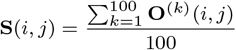

Finally, we sorted all the values in the occurrence probability matrix **S** from biggest to smallest, and denoted the *n*-th biggest value as *p_n_*. We extracted the top *αK* genes with highest occurrence probabilities to expand the prior pathways, where *α* is the parameter to control the size of expanded pathways and *K* is the total number of genes in the original pathways. The expanded pathways can be represented by the following incidence matrix:

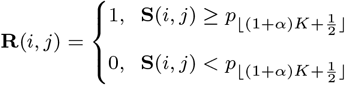

## 3 Results

### 3.1 Data Acquisition and Experimental Settings

To conduct computational experiments, we obtained 3 different survival datasets from UCSC Xena: (1) Breast Cancer Dataset (BRCA), (2) Lower Grade Glioma Dataset (LGG) and (3) Thyroid Cancer Dataset (THCA). For each cancer, we took the signaling pathways associated with the corresponding disease from KEGG DISEASE Database as the source of prior pathways, i.e. the functional modules. We only used gene expression data as the feature and the total number of genes in the original datasets is 60489. We did some preprocessing on the gene expression data. First, we transformed the read counts through log_2_(*x* + 1). Second, we selected the top variable genes of which the standard deviations among the patients were larger than 1. In this way, there were only 2005 (BRCA), 1061 (THCA) and 1126 (LGG) genes left. Third, we normalized the data into a standard normal distribution in order to overcome some problems like gradient vanishing in the neural network models. The detail information of cancer datasets and prior pathways were summarized in supplementary Table S1 and S2.

Ten-fold cross-validation was used in the two-phase training. That is, we randomly divided the samples into training set and the testing set with the ratio of 9:1. We calculated the objective function, i.e., the loss function in the training set, and simultaneously computed the evaluation metric, i.e., C-index, to monitor the performance of models in both the training set and the testing set, as shown in **Fig.** 2c. The penalty weight λ =1 in the pre-training phase and *μ* = 1 in the training phase. We adopted the Adam optimizer to train our model, in which the learning rate was set to 0.05, the number of epochs was 200, and the full batch was used. The parameter of pathway expansion *α* is set to 0.2 in the study.

**Fig. 2.**
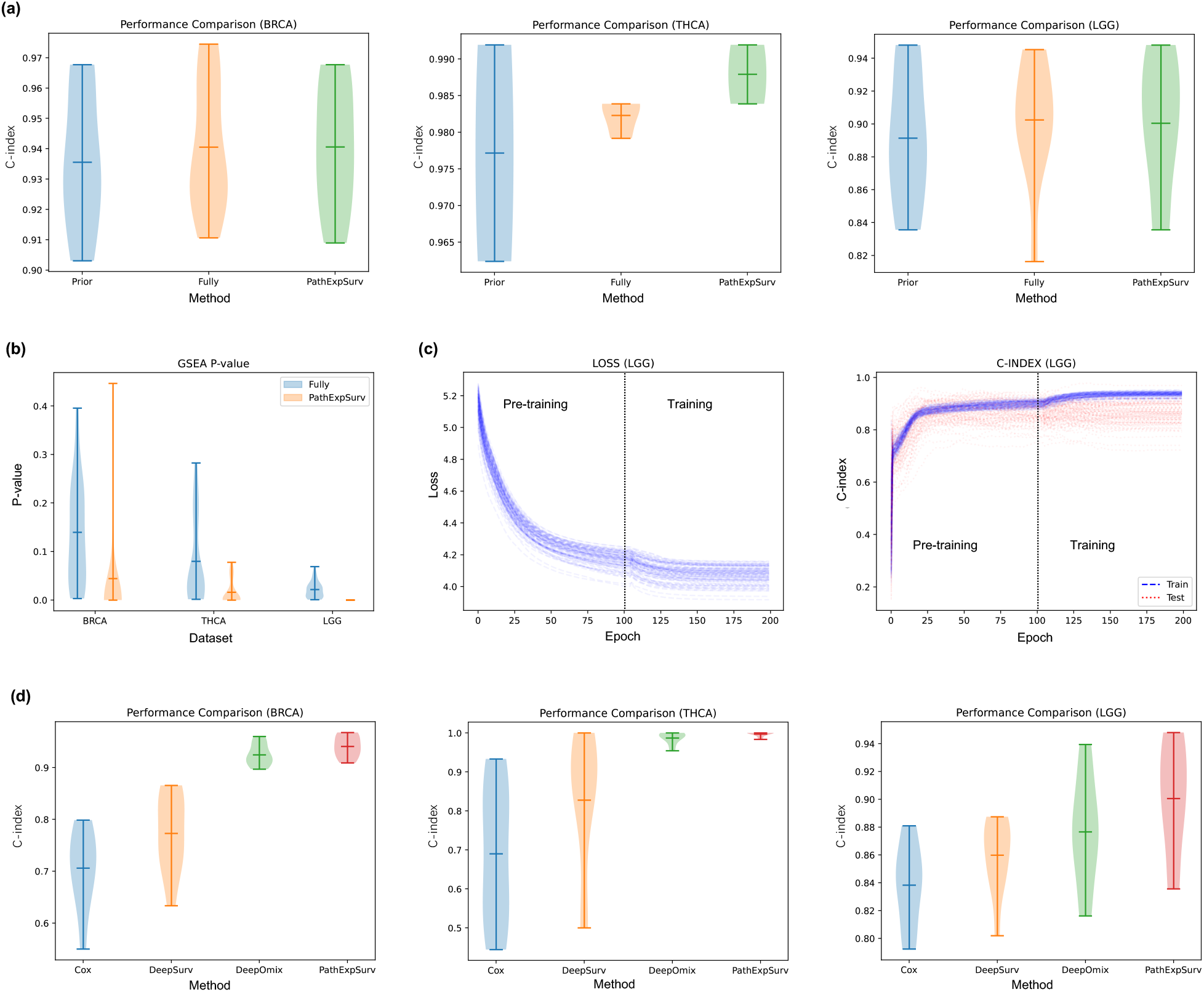
(a) Performance comparison on Prior Net, Fully-connected Net and PathExpSurv. Generally, the Fully-connected Net and PathExpSurv outperformed the Prior Net. On the THCA dataset, PathExpSurv even showed better result than the Fully-connected Net which had more learnable parameters. (b) Example of training curves of the two-phase training. The loss and C-index showed significant improvement in the training phase. (c) GSEA p-values of the ranked genes list for each pathway. The GSEA p-values of PathExpSurv are significantly smaller than those of Fully-connected Net, indicating PathExpSurv has the ability to obtain meaningful expanded pathways and the results is more interpretable. (d) Performance comparison on several methods of cancer survival analysis. The C-index results of 4 methods (Cox regression, DeepSurv, DeepOmix and PathExpSurv) are shown, and PathExpSurv had best performance among these methods.

### 3.2 Performance of Survival Analysis

We first compared the performance of PathExpSurv to two baseline models: Prior Net, Fully-connected Net. The Prior Net model only used the neural network derived from the prior pathways, and was trained using the same loss with standard deviation penalty as the pre-training phase of PathExpSurv. The Fully-connected Net model only used the fully connected neural network, and was trained using the same loss with the *L*_1_ penalty as the training phase of PathExpSurv. For fair comparison, the number of epochs of the training process was set to 200 for both Prior Net and Fully-connected Net. The training scheme of PathExpSurv can be regarded as a mixture of two baseline models, which includes 100 epochs pre-training with Prior Net and another 100 epochs training with Fully-connected Net. We performed 10-fold cross validation and the results were showed in **Fig.** 2a. As expected, the Fully-connected Net and PathExpSurv outperformed the Prior Net. On the THCA dataset, PathExpSurv even showed better result than the Fully-connected Net which had more learnable parameters.

We further investigated and compared the interpretability of PathExpSurv with the Fully-connected Net. We extracted the ranked gene list for each pathway from the weight matrix **W**1, and performed Gene Set Enrichment Analysis (GSEA) to test whether the ranked gene list is closely associated with some functional term. The p-values of the top enriched term for each pathway were shown in **Fig.** 2b. The GSEA p-values of PathExpSurv were significantly smaller than those of Fully-connected Net, indicating that PathExpSurv had the tendency to discover some genes which were closely related with each other and was more explainable than Fully-connected Net. Together with the results in **Fig.** 2a, we can conclude that the Prior Net has good interpretability but its performance might be limited, while the Fully-connected Net has higher performance but its interpretability might be poor. PathExpSurv could balance the performance and the interpretability well.

For accurately evaluating the roles of pre-training phase and training phase, we performed two-phase training scheme for 100 random experiments and computed the means and standard deviations of the results. Table 1 displayed the results of these two phases. **Fig.** 2c showed the training curve on LGG, and the training curves of other datasets were shown in supplementary **Fig.** S1. We found that the optimal C-indices of training phase were mostly better than those of pre-training phase, which meant that the training of pre-training phase learned more useful information beyond the prior pathway modules.

**Table 1.** Means and standard deviations of C-index in pre-training and training phase. The best C-index in pre-training phase and training phase is marked in bold.

| Dataset | Samples | Pre-training Phase | Training Phase |
| --- | --- | --- | --- |
| BRCA | Training set | 0.93660 ± 0.00425 | **0.95611** ± 0.00385 |
| | Testing set | 0.92812 ± 0.02134 | **0.93020** ± 0.01934 |
| THCA | Training set | 0.98640 ± 0.00315 | **0.98880** ± 0.00303 |
| | Testing set | 0.98481 ± 0.03253 | **0.98994** ± 0.01414 |
| LGG | Training set | 0.90174 ± 0.01049 | **0.93691** ± 0.00783 |
| | Testing set | **0.88602** ± 0.03614 | 0.88339 ± 0.03782 |

Finally, in order to evaluate the performance of PathExpSurv using the state-of-the-art methods, we performed 10-fold cross validation and compared the final C-index values in the testing set for each method. The performance of PathExpSurv was compared with three typical survival analysis methods: the Cox proportional hazards model (Cox, 1972), DeepSurv (Katzman *et al*., 2018), and DeepOmix (Zhao *et al*., 2021). As shown in **Fig.** 2d, we found that PathExpSurv had best performance among these methods. It is worthy to note that, the poor performance of DeepSurv is partially attributed to the over-fitting in the training dataset, while the prior information utilized in PathExpSurv and DeepOmix can help them to avoid the over-fitting.

### 3.3 Pathway Expansion

Applying the pathway expansion procedure, we identified the supplement genes of each prior pathway for each dataset, as shown in Table 2. In each disease dataset, the number of supplement genes is 20% of the total size of the original pathways. The occurrence probabilities of these supplement genes were exhibited in **Fig.** 3a, most of which are larger than 0.6, indicating these genes can be reliably identified. On the one hand, these supplement genes are significantly related to the corresponding pathway, as validated by the enrichment analysis and the recoverability testing in this section. On the other hand, these supplement genes are also closely associated with the corresponding disease, which will be demonstrated in next section.

**Fig. 3.**
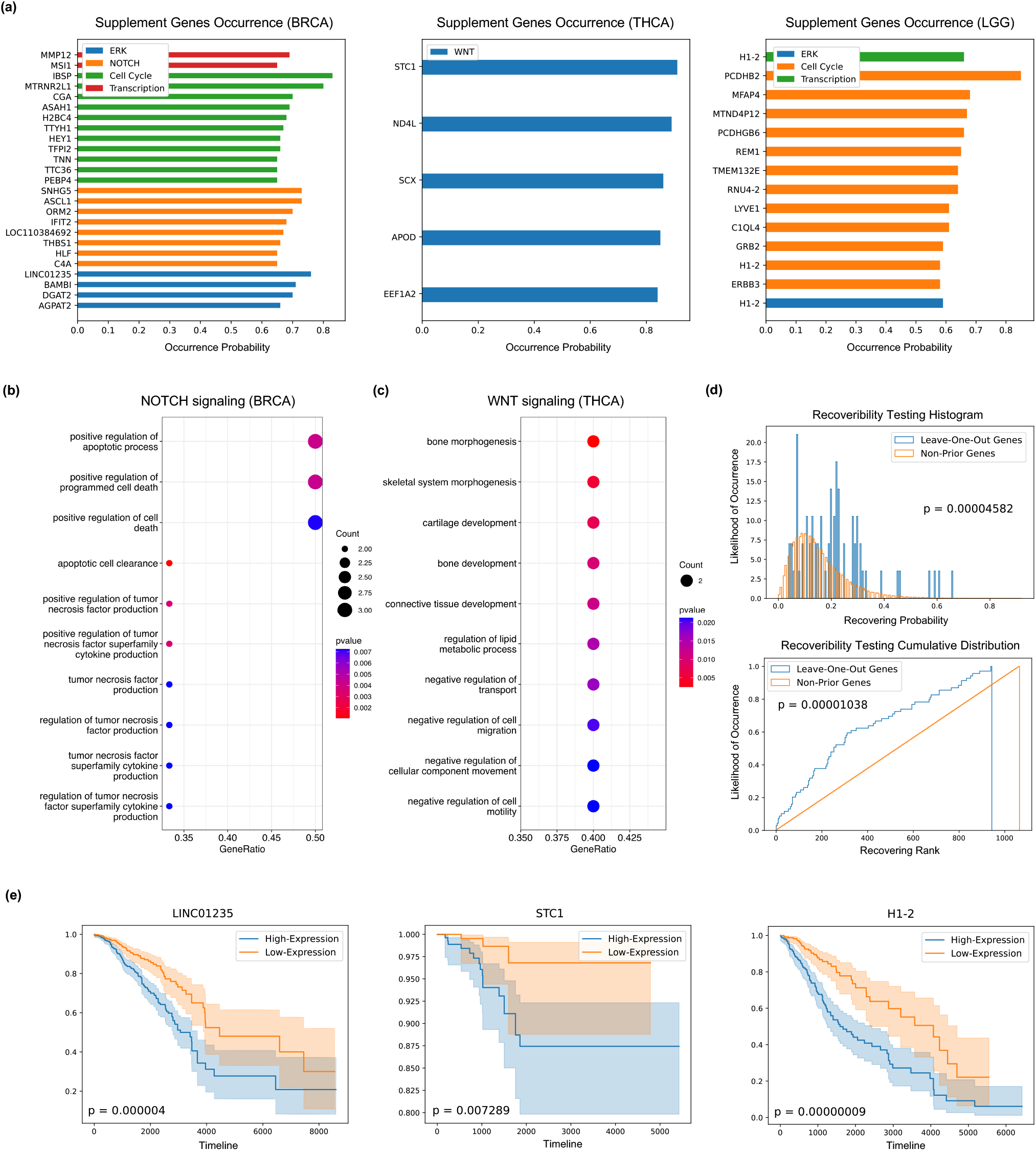
(a) Occurrence probability of the supplement genes. (b) GO term enrichment analysis result of the supplement genes of NOTCH signaling pathway for BRCA, and (c) WNT signaling pathway for THCA. (d) Comparison of the recovering probability (top) and rank (bottom) distributions of leave-one-out genes and non-prior genes. The p-values of Kolmogorov-Smirnov test are shown in the figure. (e) Kaplan-Meier curves of single-gene survival analysis for three most significantly different genes (*p* < 0.05).

**Table 2.**
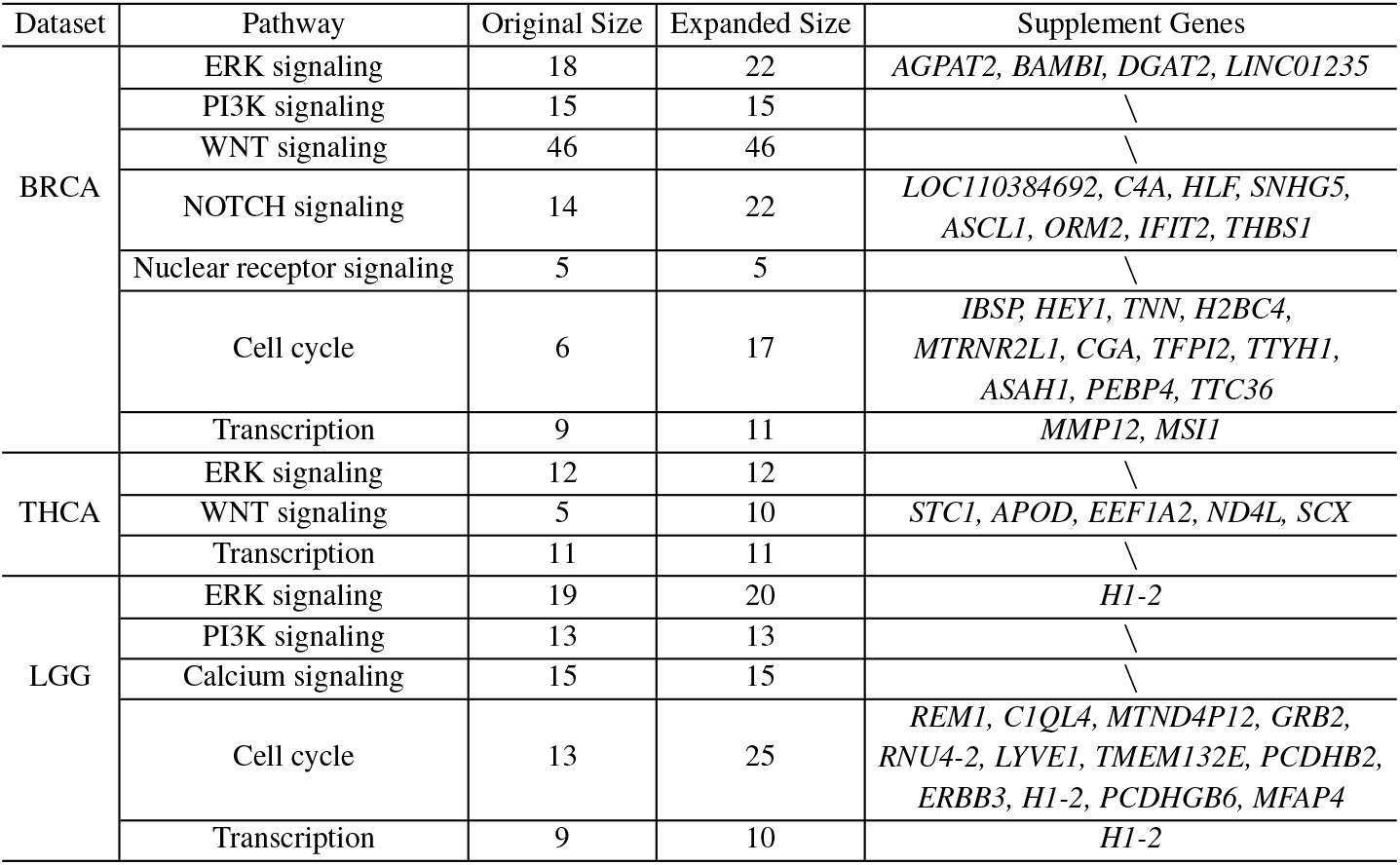
List of prior pathways and supplement genes.

We performed Gene Ontology (GO) term enrichment analysis on the supplement genes of each pathway, so as to discover the relationships between original pathway and expanded pathway. As shown in supplementary **Fig.** S3 and Table S5, the supplement genes of ERK signaling pathway for BRCA are enriched in *glycerolipid biosynthetic process* (*p* = 0.000720304) and *glycerolipid metabolic process* (*p* = 0.002490982), which are closely related to ERK signaling (Kim *et al*., 2020). The supplement genes of NOTCH signaling pathway for BRCA are enriched in *positive regulation of tumor necrosis factor production* (*p* = 0.003496794) and *positive regulation of tumor necrosis factor superfamily cytokine production* (*p* = 0.003496794), as shown in **Fig.** 3b and supplementary Table S6. Fernandez *et al*. (2008) showed that tumor necrosis factor-α modulate NOTCH signaling in the bone marrow microenvironment during inflammation. The supplement genes of WNT signaling pathway for THCA are enriched in *bone morphogenesis* (*p* = 0.00252103) and *skeletal system morphogenesis* (*p* = 0.005187936), as shown in **Fig.** 3c and supplementary Table S7. WNT signaling activates bone morphogenetic protein 2 expression (Zhang *et al*., 2013).

We also conduct a simulation experiment, named recoverability testing, to test whether PathExpSurv could recover the meaningful genes closely related to the prior pathway. We adopted the leave-one-out cross-validation strategy. Each time we removed one gene from the prior pathway and applied PathExpSurv 100 times to see how many times the leave-one-out gene can be recovered. The recovering probabilities of leave-one-out genes are compared with non-prior genes. The two-sample Kolmogorov-Smirnov test reveals that there is a significant difference between the recovering probability (rank) distributions of leave-one-out genes and non-prior genes (**Fig.** 3d). The discrepancy of the two distributions showed that the leave-one-out genes were more likely to be recovered, which might indicate that PathExpSurv had the ability to identify the genes significantly related to the corresponding pathway.

### 3.4 Disease Gene Discovery

The supplement genes are identified because they could improve the performance of survival analysis, therefore it is expected that these genes are closely associated with the corresponding disease. We searched literatures and found some promising evidence. Therefore, these genes could be further investigated and potentially used as the additional important indicators for the disease.

For breast cancer, Wang *et al*. (2015) showed the close relationship between the expression of *BAMBI* and the proliferation and migration of breast cancer. The high expression of *LINC01235* was associated with poor prognosis of breast cancer patients (Li *et al*., 2021). *IFIT2* was considered a tumor suppressor in breast cancer (Zhang *et al*., 2020), as it had been identified to inhibit cancer cell growth and migration, and promoted cell apoptosis. Chi *et al*. (2019) demonstrated that small nucleolar RNA host gene 5 (*SNHG5*) promoted breast cancer cell proliferation both in vitro and in vivo. HLF regulates ferroptosis, development and chemoresistance of triple-negative breast cancer by activating tumor cell-macrophage crosstalk (Li *et al*., 2022). The expression of *THBS1* in breast cancer was associated with poor metastasis-free survival (Yee *et al*., 2009). Knockdown of *PEBP4* inhibited breast cancer cell proliferation in vitro and tumor growth in vivo (Wang *et al*., 2017). The abnormal expression of the *IBSP* gene was closely related to bone metastasis, increased malignant risk and the poor prognosis of breast cancer (Wang *et al*., 2019). *TFPI2* was down-regulated in breast cancer tissues and cell lines, and was associated with poor prognosis of patients diagnosed with breast cancer (Zhao *et al*., 2020). Zhou *et al*. (2022) found that increased *CGA* expression was significantly associated with a poor prognosis in patients with breast cancer. *H2BC4* was overexpressed in breast cancer (Mohamed *et al*., 2021). *MSI1* was a negative prognostic indicator of breast cancer patient survival, and was indicative of tumor cells with stem cell-like characteristics (Wang *et al*., 2010).

For thyroid cancer, Hayase *et al*. (2015) demonstrated that *STC1* was highly expressed in thyroid tumor cell line and thyroid tumor tissues. The expression level of *APOD* showed significant differences in the high- and low-risk groups of differentiated thyroid cancer recurrence (Ruchong *et al*., 2021). *EEF1A2* was previously suggested as driver of tumor progression and potential biomarker.

For lower grade glioma, *ERBB3* showed marked underexpression in most glioblastomas (Duhem-Tonnelle *et al*., 2010). *GRB2* was largely involved in multiple tumor malignancies (Ijaz *et al*., 2017). Yang *et al*. (2019) indicated that *MFAP4* could be used as novel biomarker for developing therapies against human cancers.

We also performed the single-gene survival analysis to validate the significance of the newly found genes. For one specific gene, we divided the dataset into two groups: high expression group contained the top 50% gene expression level and low expression group contained the others. Then we ploted the Kaplan-Meier curves of the two groups, and identified the most significantly different genes (*p* < 0.05). We displayed three examples *(LINC01235, STC1, H1-2)* in **Fig.** 3e, while the complete curves of all the significant genes were shown in supplementary **Fig.** S4. For BRCA, we identified key genes: *LINC01235, TTC36, H2BC4, THBS1, AGPAT2, MMP12*. For THCA, we got *STC1, ND4L, APOD*. For LGG, we obtained *H1-2, LYVE1, MFAP4, PCDHGB6*. These genes were differentially expressed between two groups and might contribute to the performance improvement.

## 4 Conclusion and Discussion

In this paper, we proposed a novel survival analysis method, PathExpSurv, which exploited a two-phase training scheme to pre-train the biologically informed neural network and then train to make an exploration beyond the prior database. We showed that the pathway expansion approach can improve the performance of survival analysis while keep good interpretability of the model. Besides the survival analysis, the new method can also obtain valuable supplement genes which are significantly associated with the prior pathways and the diseases.

Although PathExpSurv has achieved good performance and showed great explainability, there still exist some directions to improve this model. Firstly, the genes beyond the database were selected based on the idea from LASSO in PathExpSurv, and we can also consider some attribution methods such as DeepLIFT (Shrikumar *et al*., 2017), DeepExplain (Ancona *et al*., 2017) and LIME (Ribeiro *et al*., 2016). Secondly, PathExpSurv only employed a 3-layer neural network, and more sophisticated architecture might further improve the performance and interpretability. Finally, while the training scheme of PathExpSurv consisted of two phases, we can design a more complex training way to adjust the pathways step by step. Furthermore, PathExpSurv could be regarded as a high-level framework which might be applied to all kinds of prediction tasks.

## Funding

This work was supported by the National Key Research and Development Program of China [2020YFA0712402] and National Natural Science Foundation of China [12231018].

## Notes

### Competing Interest Statement

The authors have declared no competing interest.

### Summary of Updates

Correct the error of URL

