## Supplementary Materials for "PathExpSurv: Pathway Expansion for Explainable Survival Analysis and Disease Gene Discovery"

[1]IAM, MADIS, NCMIS, Academy of Mathematics and Systems Science, Chinese Academy of Sciences, Beijing 100190, China, [2]School of Mathematical Sciences, University of Chinese Academy of Sciences, Beijing 100049, China, [3]Computational Biology Program, Oregon Health & Science University, Portland, OR 97239, USA, [4]Department of Molecular Microbiology and Immunology, Oregon Health & Science University, Portland, OR 97239, USA.

### 1 Foundation of Survival Time Analysis

#### 1.1 Notations of Common Terms in Survival Analysis

Survival analysis is used to handle the survival data which is right-censored (i.e. some patients may leave the study or the study ends before an event occurs).

The compositions of survival data are: a patient's observed covariates $\mathbf{x}$, a survival time $T$, and an event indicator $\delta \in \{0, 1\}$.

The survival function is denoted as

$$S(t) = P(T > t),$$

which signifies the probability that the failure time is latter than $t$.

The hazard function is defined as:

$$h(t) = \lim_{\Delta t \to 0} \frac{P(t \leq T < t + \Delta t \mid T \geq t)}{\Delta t}$$

It is the probability an individual will not survive for an additional time $\Delta t$, given they have already survived up to time $t$.

#### 1.2 Cox Proportional Hazards Model

The Cox proportional hazards model is a common method for modeling an individual's survival given the feature $\mathbf{x}$. The model assumes that the hazard function is composed of two non-negative functions: a baseline hazard function, $h_0(t)$, and a risk score, $r(\mathbf{x}) = \exp(\mathbf{x}^T \beta)$, defined as the effect of an individual's obeserved covariates on the baseline hazard. Then the hazard function is assumed to have the form:

$$h(t \mid x) = h_0(t) \exp\left(\mathbf{x}^T \beta\right)$$

The likelihood of the event to be observed occurring for subject $i$ at time $T_i$ can be written as:

$$L_i(\beta) = \frac{h(T_i \mid \mathbf{x}_i)}{\sum_{j: T_j \geq T_i} h(T_j \mid \mathbf{x}_j)} = \frac{h_0(T_i) \exp\left(\mathbf{x}_i^\top \beta\right)}{\sum_{j: T_j \geq T_i} h_0(T_i) \exp\left(\mathbf{x}_j^\top \beta\right)} = \frac{\exp\left(\mathbf{x}_i^\top \beta\right)}{\sum_{j: T_j \geq T_i} \exp\left(\mathbf{x}_j^\top \beta\right)}$$

The Cox log partial likelihood function is defined as:

$$L_c(\beta) = \log\left(\prod_{i: \delta_i = 1} L_i(\beta)\right) = \sum_{i=1}^{n} \delta_i \left[\mathbf{x}_i^\top \beta - \log\left(\sum_{j: T_j \geq T_i} \exp\left(\mathbf{x}_j^\top \beta\right)\right)\right]$$

The commonly used metric in survival analysis is concordance index (C-index):

$$\textbf{C-index} = \frac{\sum_{i,j} 1_{T_j < T_i} \cdot 1_{r(\mathbf{x}_j) > r(\mathbf{x}_i)} \cdot \delta_j}{\sum_{i,j} 1_{T_j < T_i} \cdot \delta_j}$$

### 2 Details of Data

#### 2.1 Cancer Datasets

For data acquisition, we obtained 3 different survival datasets from UCSC Xena: (1) Breast Cancer Dataset (BRCA), (2) Lower Grade Glioma Dataset (LGG), (3) Thyroid Cancer Dataset (THCA). Table S1 shows the number of samples and selected genes of 3 different datasets. For each cancer, we only used gene expression data as the feature and the total number of genes was 60489. Then we did some preprocessing for the gene expression data. First, we transformed the read counts through $\log_2(x + 1)$. Second, we selected the top variable genes of which the standard deviations among the patients were larger than 1. In this way there were only 2005 (BRCA), 1061 (THCA) and 1126 (LGG) genes left. Third, we normalized the data into a standard normal distribution in order to overcome some problems like gradient vanishing in the deep learning models.

Table S1: Datasets basic information.

| | (num-samples, num-genes) |
|---|---|
| BRCA | (1194, 2005) |
| THCA | (567,1061) |
| LGG | (524,1126) |

#### 2.2 KEGG Signaling Pathways

We took the prior pathways as the functional modules and the source of the prior signaling pathways was KEGG DISEASE Database. The details of the composition of the pathways are showed in Table S2 .

Table S2: Compositions of the prior pathways.

| Disease | Pathway | Composition |
|---|---|---|
| BRCA | ERK signaling | *HRAS/BRAF/EGFR/GRB2/KRAS/RAF1/MAP2K1/ MAP2K2/ERBB2/ARAF/EGF/FGFR1/FGFR2/MAPK1/ MAPK3/NRAS/SOS1/SOS2* |
| | PI3K signaling | *PTEN/EGFR/PIK3CA/PIK3CB/MTOR/PIK3CD/AKT1/AKT2/ AKT3/ERBB2/FGFR1/RPS6KB1/RPS6KB2/FGFR2/EGF* |
| | WNT signaling | *MYC/FZD1/FZD4/FZD7/WNT5B/FZD6/FZD8/FZD9/TCF7/ TCF7L2/FRAT1/FZD3/WNT1/WNT2/WNT3/WNT5A/WNT6/ WNT3A/WNT7A/WNT7B/WNT8A/WNT10B/WNT8B/WNT2B/ WNT9A/WNT9B/WNT16/DVL1/DVL2/DVL3/APC/WNT10A/ LRP6/LRP5/FZD10/CCND1/WNT4/CTNNB1/FZD5/ FZD2/LEF1/FRAT2/TCF7L1/GSK3B/AXIN1/AXIN2* |
| | NOTCH signaling | *DLL1/JAG2/HEY2/HEY1/DLL4/HES5/HEYL/ ERBB2/HES1/NOTCH1/FLT4/JAG1/NOTCH4/ DLL3* |
| | Nuclear receptor signaling | *MYC/NCOA1/NCOA3/CCND1/ESR1* |
| | Cell cycle | *RB1/E2F1/E2F2/E2F3/CCND1/CDK4* |
| | Transcription | *GADD45G/CDKN1A/BAK1/POLK/BAX/ GADD45B/DDB2/GADD45A/TP53* |
| THCA | ERK signaling | *NRAS/ARAF/BRAF/RAF1/MAP2K1/MAP2K2/ MAPK1/MAPK3/RET/HRAS/KRAS/CCND1* |
| | WNT signaling | *TCF7L1/TCF7L2/LEF1/MYC/CCND1* |
| | Transcription | *GADD45B/GADD45G/BAX/BAK1/DDB2/ POLK/PAX8/RXRA/RXRB/RXRG/PPARG* |
| LGG | ERK signaling | *KRAS/RAF1/NRAS/BRAF/PDGFA/PDGFB/EGFR/ PDGFRA/PDGFRB/HRAS/GRB2/CCND1/MAPK1/ MAPK3/MAP2K1/MAP2K2/ARAF/SOS1/SOS2* |
| | PI3K signaling | *PTEN/EGFR/PDGFRA/PIK3CA/PIK3CB/MTOR/ PIK3CD/AKT1/AKT2/AKT3/RPS6KB1/RPS6KB2/BAD* |
| | Calcium signaling | *CALM1/CAMK1/EGFR/CALM2/PDGFRA/CALM3/ CAMK4/CAMK2A/CAMK2B/CAMK2D/CAMK2G/ CAMK1G/PLCG1/PLCG2/CAMK1D* |
| | Cell cycle | *CCND3/MDM2/CDKN1A/CDKN2A/RB1/E2F1/E2F2/ E2F3/CCND1/TP53/CDK4/CDK6/CCND2* |
| | Transcription | *GADD45G/CDKN1A/BAK1/POLK/BAX/ GADD45B/DDB2/GADD45A/TP53* |

### 3 Experiment Results

#### 3.1 Two-phase Training

Table S3 displayed the exact results of these two phases, and Figure S1 showed the training curves. We found that the optimal c-indices of training phase were mostly better than those of pre-training phase, which means that the training of pre-training phase learned more useful information beyond the prior pathways modules.

Table S3: Results of pre-training & training phase.

| Disease | Item | Pre-training Phase | Training Phase |
|---|---|---|---|
| BRCA | Loss | 4.76112 ± 0.03988 | **4.69065** ± 0.04615 |
| | C-index (train) | 0.93660 ± 0.00425 | **0.95611** ± 0.00385 |
| | C-index (test) | 0.92812 ± 0.02134 | **0.93020** ± 0.01934 |
| THCA | Loss | 3.58619 ± 0.16515 | **3.51818** ± 0.16522 |
| | C-index (train) | 0.98640 ± 0.00315 | **0.98880** ± 0.00303 |
| | C-index (test) | 0.98481 ± 0.03253 | **0.98994** ± 0.01414 |
| LGG | Loss | 4.16411 ± 0.05032 | **4.07388** ± 0.05227 |
| | C-index (train) | 0.90174 ± 0.01049 | **0.93691** ± 0.00783 |
| | C-index (test) | **0.88602** ± 0.03614 | 0.88339 ± 0.03782 |

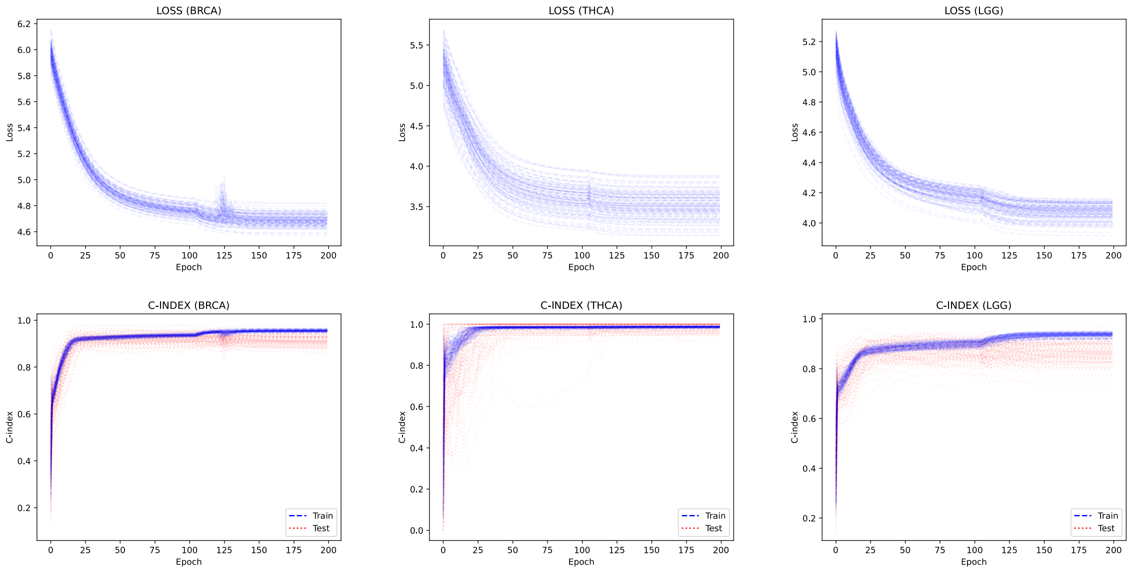

Figure S1: Training curves of pre-training and training phase. The model achieves better performance in training phase (2nd 100 epoches) than that in pre-training phase (1st 100 epoches).

#### 3.2 Retraining

Table S4 and Figure S2 compared the results of model with original pathways mask and expanded pathways mask. The results showed that the expanded pathways achieved better performance than original pathways. We found that the expanded pathways outperformed significantly original modules at epoch 5, which indicated that the newly learned meta-pathways could help researchers to predict more quickly. We noticed that the C-index on the training set of expanded pathways was worse than that of original pathways, but the result was converse on testing set, it might demonstrate that the meta-pathways overcome the over-fitting problem in some way.

Table S4: Comparison of original pathways and expanded pathways.

| Disease | Item | Original | Expanded |
|---|---|---|---|
| BRCA | Loss | 4.76112 ± 0.03988 | **4.78944** ± 0.03473 |
| | C-index (Train Optimal) | **0.93660** ± 0.00425 | 0.92501 ± 0.00466 |
| | C-index (Train Epoch 5) | 0.73742 ± 0.01801 | **0.78574** ± 0.03140 |
| | C-index (Test Optimal) | 0.92812 ± 0.02134 | **0.92828** ± 0.02094 |
| | C-index (Test Epoch 5) | 0.71595 ± 0.07056 | **0.77376** ± 0.06358 |
| THCA | Loss | 3.58619 ± 0.16515 | **3.55375** ± 0.16606 |
| | C-index (Train Optimal) | **0.98640** ± 0.00315 | 0.98594 ± 0.00396 |
| | C-index (Train Epoch 5) | 0.86103 ± 0.04687 | **0.91480** ± 0.03014 |
| | C-index (Test Optimal) | 0.98481 ± 0.03253 | **0.98906** ± 0.01905 |
| | C-index (Test Epoch 5) | 0.83347 ± 0.15593 | **0.90224** ± 0.10201 |
| LGG | Loss | 4.16411 ± 0.05032 | **4.14754** ± 0.04814 |
| | C-index (Train Optimal) | 0.90174 ± 0.01049 | **0.90427** ± 0.00966 |
| | C-index (Train Epoch 5) | 0.74366 ± 0.02394 | **0.75932** ± 0.02198 |
| | C-index (Test Optimal) | **0.88602** ± 0.03614 | 0.87816 ± 0.04556 |
| | C-index (Test Epoch 5) | 0.74234 ± 0.06984 | **0.75472** ± 0.06494 |

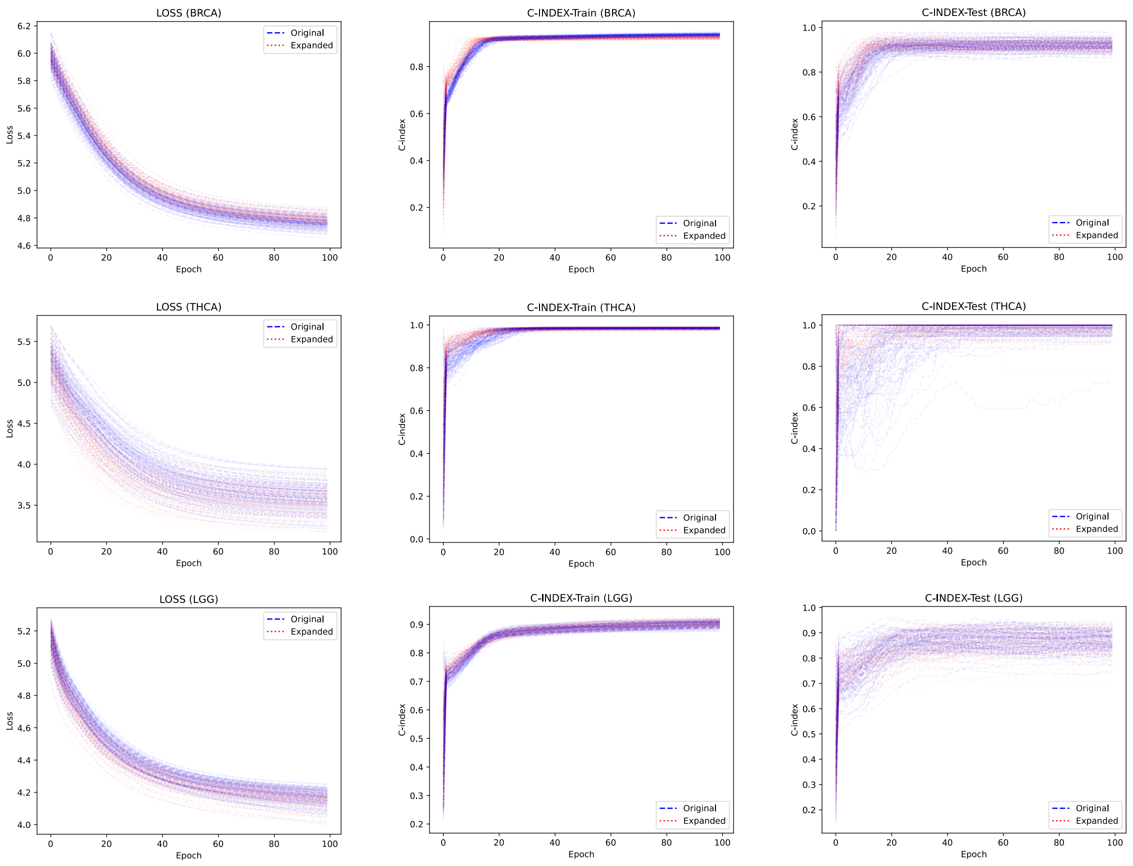

Figure S2: Training curves of retraining. The expanded-pathways model can achieve better performance quickly and finally outperform original-pathways model.

### 4 Analysis of Expanded Pathways

#### 4.1 Gene Ontology (GO) Term Enrichment Analysis

We performed Gene Ontology (GO) term enrichment analysis for the supplement genes of every pathway, so as to discover some relationships between original pathways and expanded pathways. The results are shown in Figure S3, and the details for the supplement genes of ERK signaling pathway (BRCA), NOTCH signaling pathway (BRCA) and WNT signaling pathway (THCA) are shown in Tables S5, S6 and S7, respectively.

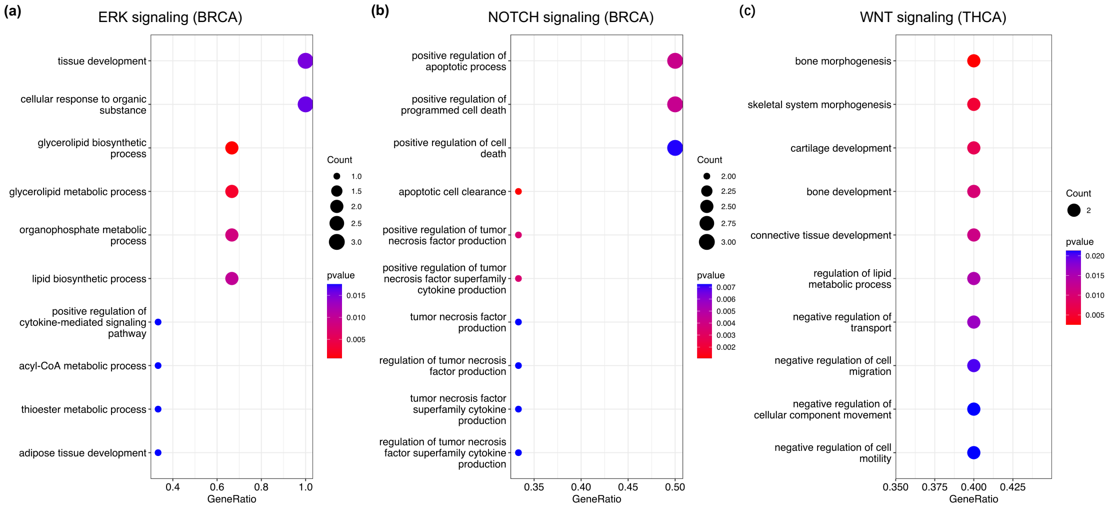

Figure S3: Go Enrichment Analysis

Table S5: GO Enrichment Analysis (ERK signaling - BRCA)

| Description | GeneRatio | BgRatio | pvalue | geneID |
|---|---|---|---|---|
| glycerolipid biosynthetic process | 2/3 | 27/1702 | 0.000720304 | 84649/10555 |
| glycerolipid metabolic process | 2/3 | 50/1702 | 0.002490982 | 84649/10555 |
| organophosphate metabolic process | 2/3 | 92/1702 | 0.008369153 | 84649/10555 |
| lipid biosynthetic process | 2/3 | 101/1702 | 0.010059627 | 84649/10555 |
| tissue development | 3/3 | 422/1702 | 0.015161112 | 84649/10555/25805 |
| cellular response to organic substance | 3/3 | 428/1702 | 0.015818615 | 84649/10555/25805 |
| positive regulation of cytokine-mediated signaling pathway | 1/3 | 10/1702 | 0.017533207 | 10555 |
| acyl-CoA metabolic process | 1/3 | 10/1702 | 0.017533207 | 84649 |
| thioester metabolic process | 1/3 | 10/1702 | 0.017533207 | 84649 |
| adipose tissue development | 1/3 | 10/1702 | 0.017533207 | 84649 |

Table S6: GO Enrichment Analysis (NOTCH signaling - BRCA)

| Description | GeneRatio | BgRatio | pvalue | geneID |
|---|---|---|---|---|
| apoptotic cell clearance | 2/6 | 15/1702 | 0.001066033 | 720/7057 |
| positive regulation of tumor necrosis factor production | 2/6 | 27/1702 | 0.003496794 | 7057/5005 |
| positive regulation of tumor necrosis factor superfamily cytokine production | 2/6 | 27/1702 | 0.003496794 | 7057/5005 |
| positive regulation of apoptotic process | 3/6 | 107/1702 | 0.004204728 | 7057/3433/429 |
| positive regulation of programmed cell death | 3/6 | 108/1702 | 0.004318901 | 7057/3433/429 |
| positive regulation of cell death | 3/6 | 129/1702 | 0.007181695 | 7057/3433/429 |
| tumor necrosis factor production | 2/6 | 39/1702 | 0.00724334 | 7057/5005 |
| regulation of tumor necrosis factor production | 2/6 | 39/1702 | 0.00724334 | 7057/5005 |
| tumor necrosis factor superfamily cytokine production | 2/6 | 39/1702 | 0.00724334 | 7057/5005 |
| regulation of tumor necrosis factor superfamily cytokine production | 2/6 | 39/1702 | 0.00724334 | 7057/5005 |

Table S7: GO Enrichment Analysis (WNT signaling - THCA)

| Description | GeneRatio | BgRatio | pvalue | geneID |
|---|---|---|---|---|
| bone morphogenesis | 2/5 | 14/838 | 0.00252103 | 642658/6781 |
| skeletal system morphogenesis | 2/5 | 20/838 | 0.005187936 | 642658/6781 |
| cartilage development | 2/5 | 24/838 | 0.007463464 | 642658/6781 |
| bone development | 2/5 | 28/838 | 0.010122866 | 642658/6781 |
| connective tissue development | 2/5 | 30/838 | 0.011592773 | 642658/6781 |
| regulation of lipid metabolic process | 2/5 | 34/838 | 0.014805605 | 347/1917 |
| negative regulation of transport | 2/5 | 36/838 | 0.016545606 | 347/6781 |
| negative regulation of cell migration | 2/5 | 40/838 | 0.020285551 | 347/6781 |
| negative regulation of cellular component movement | 2/5 | 41/838 | 0.021273649 | 347/6781 |
| negative regulation of cell motility | 2/5 | 41/838 | 0.021273649 | 347/6781 |

#### 4.2 Single-Gene Survival Analysis

We performed the single-gene survival analysis to validate the significance of the supplement genes in each disease. For one specific gene, we divided the dataset into two groups: high expression group contained the top 50% gene expression level and low expression group contained the others. Then we ploted the Kaplan-Meier curves of the two groups , and identified the most significantly different genes ($p < 0.05$). We displayed the Kaplan-Meier curves in Figure S4. For BRCA, we identified key genes: *LINC01235, TTC36, H2BC4, THBS1, AGPAT2, MMP12*. For THCA, we got *STC1, ND4L, APOD*. For LGG, we obtained *H1-2, LYVE1, MFAP4, PCDHGB6*. These genes were differentially expressed between two groups and might contribute to the performance improvement.

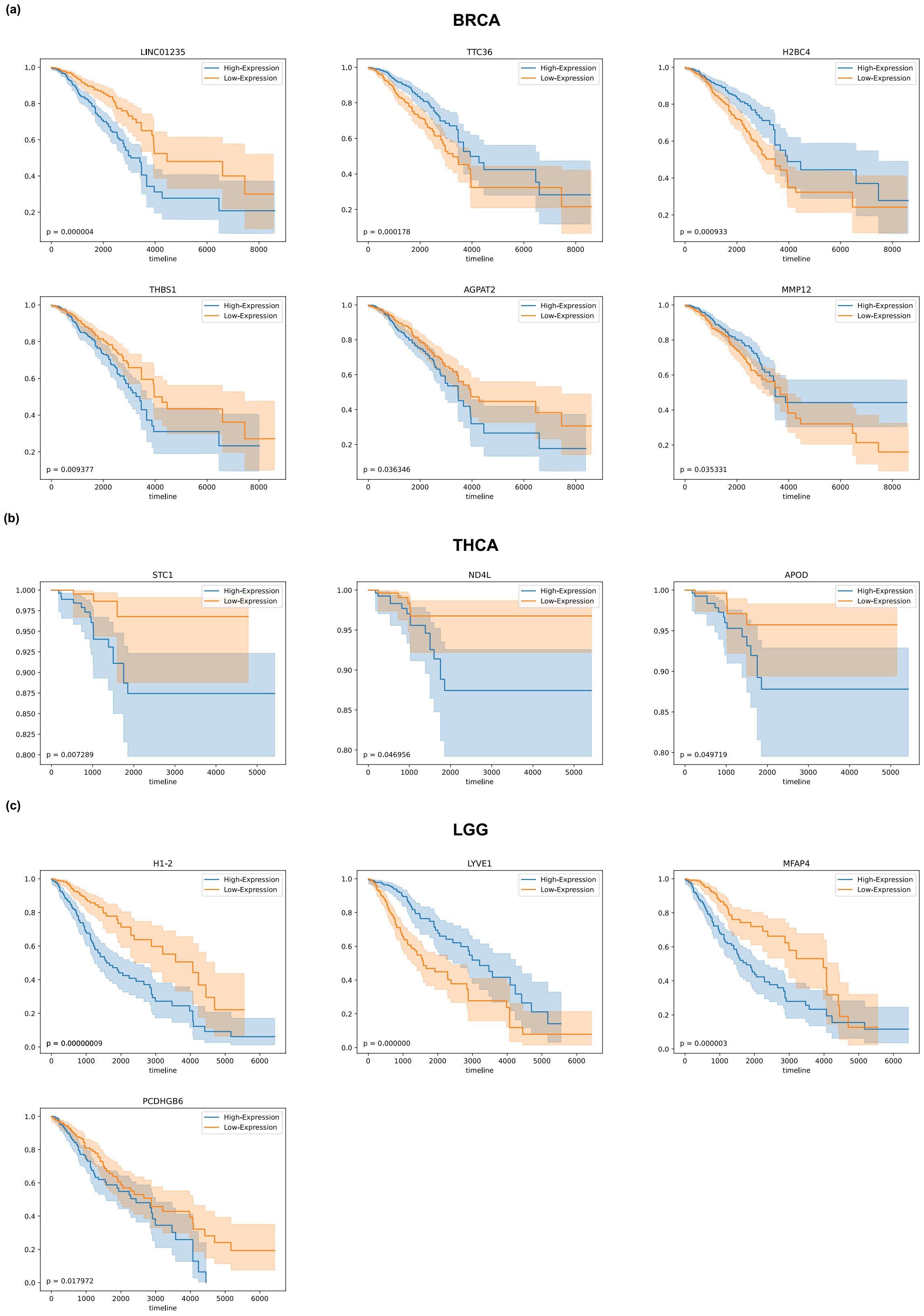

Figure S4: Kaplan Meier curves
